# Natural variation of the cardiac transcriptome in humans

**DOI:** 10.1101/2020.10.06.328591

**Authors:** Tatiana Domitrovic, Mariana H. Moreira, Rodolfo L. Carneiro, Marcelo Ribeiro-Alves, Fernando L. Palhano

## Abstract

We investigated the gene-expression variation among humans by analyzing previously published mRNA-seq and ribosome footprint profiling (RP) of heart left-ventricles from healthy donors. We ranked the genes according to their coefficient of variation values and found that the top 5% most variable genes had special features compared to the rest of the genome, such as lower mRNA levels and shorter half-lives coupled to increased translation efficiency. We observed that these genes are mostly involved with immune response and have a pleiotropic effect on disease phenotypes, indicating that asymptomatic conditions contribute to the gene expression diversity of healthy individuals.

## INTRODUCTION

The gene expression variation across different individuals arises from a complex interplay between the environment and the cellular responses to a wide range of stimuli and, therefore, can be attributed to either genetic or non-genetic factors^1^. To pinpoint the gene expression responses linked to a specific illness, one must also understand which are the genes presenting the highest degree of expression variation across the general human population. Most of the studies comparing geneexpression levels have been done in the context of disease. These studies aimed to identify changes in cellular and molecular processes associated with the illness and/or identify expression Quantitative Trait Loci (eQTLs) or splicing QTLs (sQTLs) related to the disease state. In such studies, data on gene expression variation among non-disease individuals are not explicitly discussed. Instead, this group is treated as a control, and the inter-individual variation data are embedded in the calculation of differential gene expression. By contrast, studies dealing with phenotypic and genetic diversity in human populations directly describe the natural variation of gene expression. Such studies were levered by international projects, such as HapMap^2^ and Gtex^3,4^, designed to capture the natural genetic variance of the human population. Following this rationale, different groups showed that gene expression variation between individuals is greater than the variation among populations and mapped many polymorphic genetic variances that influence gene expression differences between individuals^5–13^. Bulk genome-wide gene expression is commonly accessed at the mRNA level by microarray or by mRNA sequencing (mRNA-seq). However, the ribosome profiling (RP) technique, which captures mRNA footprints protected by translating ribosomes^14^, now allows the direct interrogation of the translatome using mRNA-seq. However, there are few studies aimed to compare gene regulation results obtained through RNA-seq and ribosome profiling^15^ and most these references focus on the disease state.

In this work, we accessed the natural variation in gene expression using a specific data set containing RNA-seq and RP data of healthy donors’ left-ventricle heart tissue. Using a stringent criterium, we selected a group of genes with a high coefficient of variation (CV) between individuals. Next, we analyzed several parameters related to gene expression control, such as translation efficiency, mRNA and protein half-life, and protein abundance. The genes with high CV presented high translation efficiency while their mRNA levels and mRNA half-lives were shorter than the genome, suggesting an optimization for fast, transient expression. Indeed, many of these genes are involved in immune response and response to virus infection and have been identified as pleiotropic factors involved in the disease phenotype. We also directly compared the most variable genes in the RNA-seq and RP profiling datasets. Even though these techniques show a good general correlation, the genes in the high CV group obtained with each technique were markedly different. This result may arise not only from technical particularities of the RP assay but also, from the additional layers of regulation influencing the translation process.

## RESULTS AND DISCUSSION

To study cardiac mRNA expression and variation among individuals, we used previously published mRNA-seq and ribosome profiling data from fifteen different donors (Supplementary Table S1). Those individuals were used as controls in a large study with dilated cardiomyopathy (DCM) patients^16^. From the clinical perspective, the fifteen individuals presented no heart-related pathology^16^. Our first aim was to compare the Spearman correlation among the mRNA-seq data e ribosome profiling (RP) data for each individual (Fig 1AB). Ribosome profiling involves similar sequencing library preparation and data analysis used for RNA-seq. However, RP involves additional steps before sequencing, such as ribosome purification and endonucleolytic cleavage of non-protected mRNA. In this way, ribosome profiling targets only mRNA sequences protected by the ribosome during decoding by translation^17^. Usually, there is a positive strong correlation between RNA-seq and RP data^18^, but this is not necessarily true for all genes. For example, genes with efficient ribosome initiation might have a large amount of RP protected reads, even with relatively few copies of mRNA^19^. Figure 1A represents a scatter plot of mRNA-seq vs. RP data from one of the individuals. We observed a strong Spearman’s correlation coefficient (ρ = 0.80) between the two data. When the same analysis was extended to the other 14 subjects, we observed correlation coefficient values ranging from 0.47 to 0.85 (Fig 1B). Then, we analyzed how the gene expression correlates between individuals. The correlation matrix using mRNA-seq data revealed two main clusters (Fig 1C). The first cluster (B, M, and N individuals) had correlation coefficient values ranging from 0.72 to 0.89, while the second cluster (A, C, D, E, F, G, H, I J, K, L, and O individuals) grouped similar individuals with ρ values ranging from 0.91 to 0.96 (Fig 1C). We observed a different clustering pattern when the RP data was used to build the matrix. The A individual was separated from others while the N individual clustered together (Fig 1D). Notably, the RP data correlations from different individuals resulted in the broader variation of correlation coefficient values (ρ = 0.58 to 0.94) than observed with correlation analysis of the RNA-seq data (ρ = 0.72 to 0.96).

**Figure 1.**
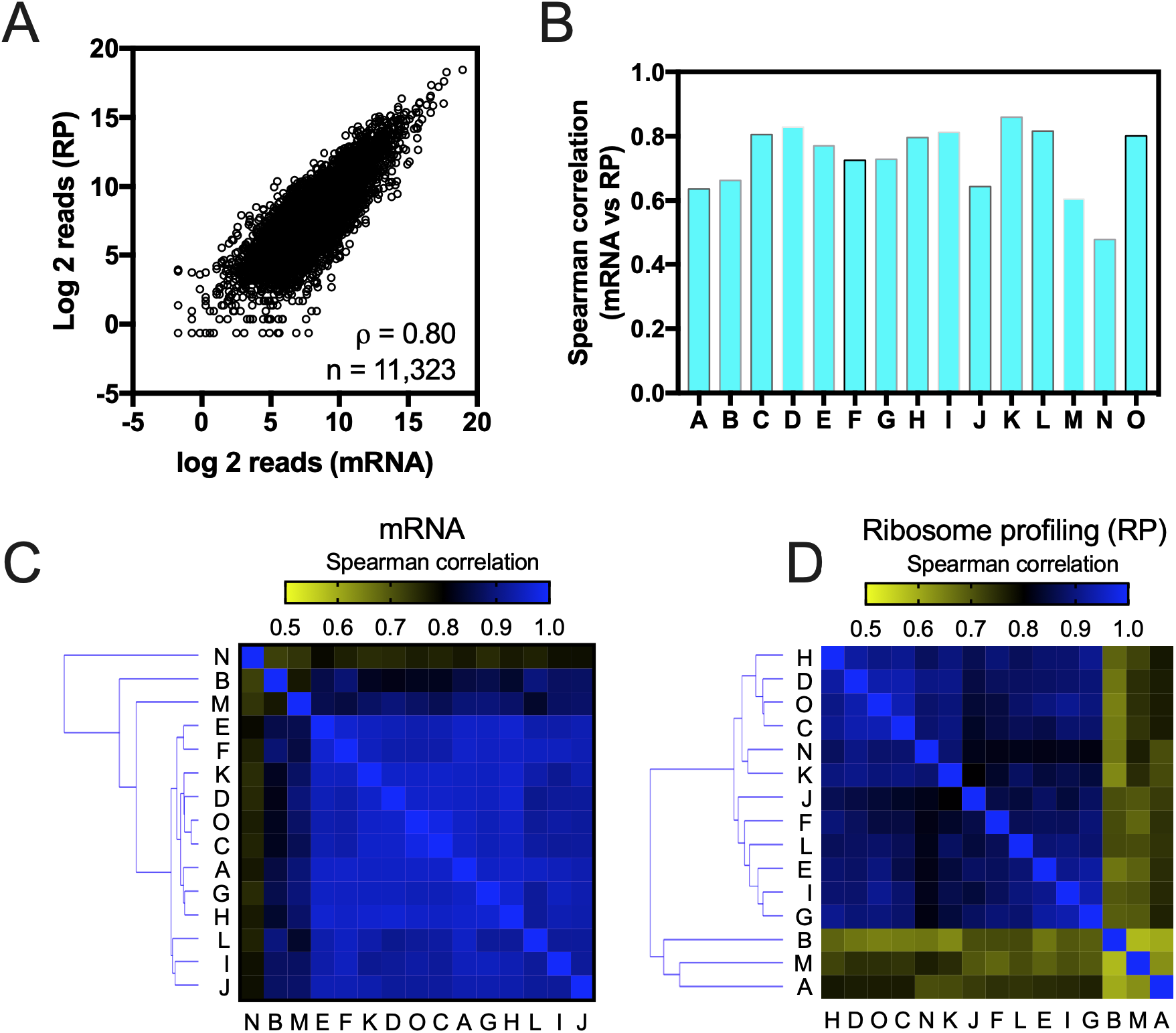
Correlation analyses between mRNA-seq data and ribosome profiling from a healthy human heart left ventricles. (A) Gene expression measured by RNA-seq compared to gene expression measured by ribosome profiling (RP). Spearman correlation (ρ) is indicated. (B) The Spearman correlation (ρ) values between RNA-seq and ribosome profiling (RP) data for each of the fifteen individuals used herein. A correlation matrix of the Spearman correlation (ρ) values obtained with RNA-seq data (C) or ribosome profiling data (D) among the individuals.

Next, we used RNA-seq data to select the group of genes that had the highest variability across healthy individuals. To make our analysis more stringent and less subject to variations from a particular subject, we excluded the three individuals that presented the lower correlation coefficients values for mRNA expression across the population (B, M, and N individuals, Fig 1C). We determined the expression variability between individuals by calculating the coefficient of variation (CV) of each gene. Figure 2A shows the CV of four representative genes. While a low variation value (CV = 0.009) was observed for the gene RTFDC1, a higher variation value (CV = 0.19) was observed for SERPINE1, comprising a 43-fold difference between the lower vs. higher values (Fig 2A). The statistical test to identify the most variable genes was based on the assumption that the top 2.5% more expressed genes are housekeeping genes of each sample. Then, the CV value of each of the other genes was compared to the mean CV value of this housekeeping group of genes. We observed that 32.2% of the genes had CV values different from the house-keeping group (p < 0.05). We focused on the top 5% genes with the highest CV (Fig 2B, 565 genes).

**Figure 2.**
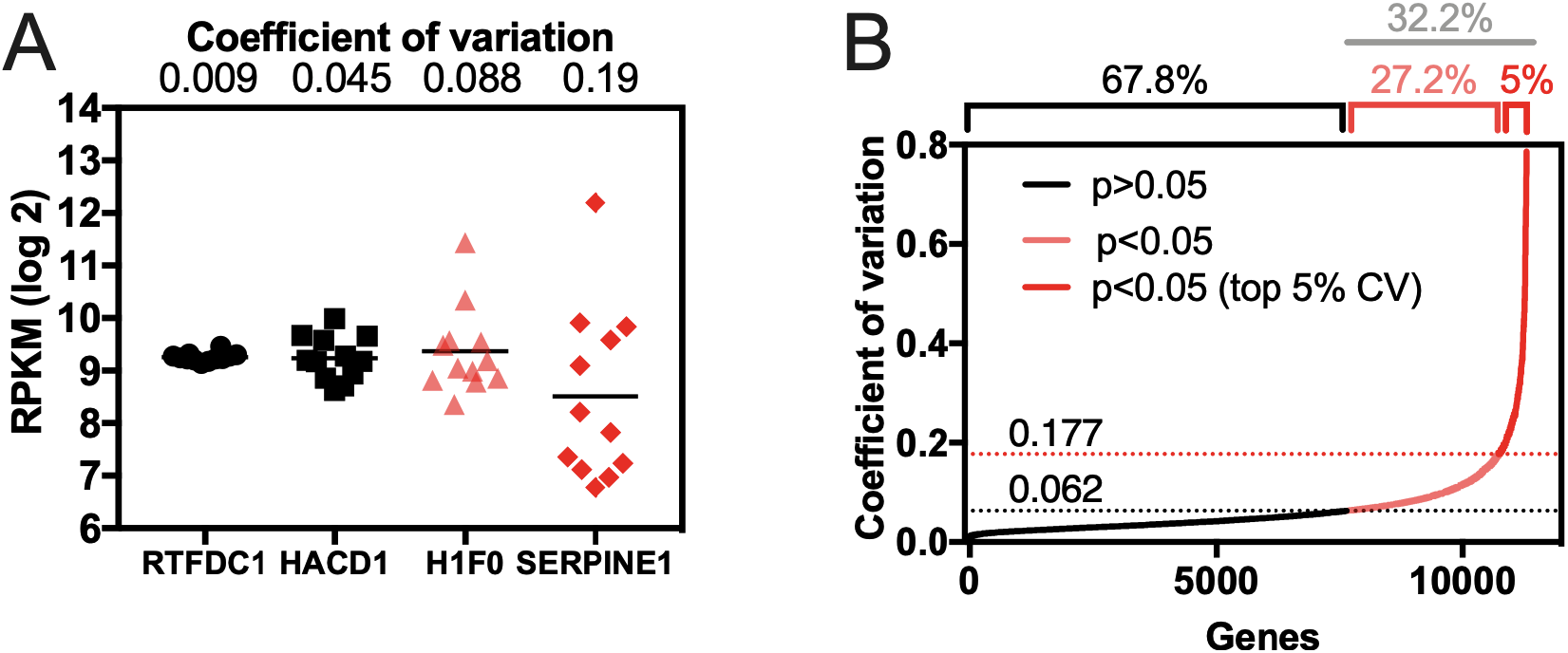
Expression variability of the heart transcriptome among individuals. (A) Example of expression variation of four genes with a variation coefficient ranging from low (0.009, RTFDC1) to high (0.19, SERPINE1). The expression of those genes was measured by RNA-seq and plotted as reads per kilobase per million (RPKM). (B) Cumulative distribution plot of the coefficient of variation values obtained from a healthy human heart left ventricles (12 individuals). Values of CV higher than 0.062 (dotted black line) were considered statistically different (p<0.05) when compared with the average CV from housekeeping genes (see details in methodology). The top 5% higher CV genes have CV values higher than 0.177 (dotted red line).

To determine if the results obtained with these 12 individuals would represent a larger human population, we analyzed RNA-seq data from 432 heart left-ventricles of different healthy donors, available through the GTex database. The mean of gene expression values for each gene of the van Heesch’s dataset and the Gtex dataset had a positive correlation coefficient of 0.84 (Fig S1A), and the CV values had a correlation coefficient of 0.51 (Fig S1B). The group of the most variable genes from the GTex (CV #2, 550 genes) contained 297 genes in common (or 54%) with van Heesch’s most variable genes (CV #1, 565 genes) (Fig S2A), and the Gene ontology analysis of the groups showed that the genes from both high CV groups were involved in similar molecular processes (Fig S2B-C), mainly inflammatory process. These results indicate that the van Heesch’s group generally reflects the results obtained with a larger RNA-seq dataset. The next objective was to determine if the genes with the most variable transcriptional levels would also have variable translational levels between individuals. For this, we repeated the methodology described above using RP data instead of RNA-seq data and obtained a group of 565 genes that were ranked as the top 5% with the highest CV values (p<0.05). When we compared the mRNA-seq high CV group, with the RP high CV group, we found only that 149 (or 26.3%) were the same. This is three-fold more than what would be expected by random (8.5%) but is 6-fold less than the intersection between Gtex and van Heesch’s RNA-seq high CV groups. This result is in agreement with van Heesch’s study, where they found that from the 2648 genes differentially expressed by RP in DCM hearts vs. control hearts, only 964 genes could have their expression levels explained by transcriptional differences. Therefore, in addition to the technical differences between mRNA and RP, a good part of the transcriptional level variance could be muffled by additional regulation layers acting on the translational level.

Next, we asked whether the group of genes with high CV values, derived from RNA-seq (van Heesch’s CV #1), would have any particular signature concerning a variety of parameters related to gene expression. For this analysis, protein abundance^20^, mRNA abundance^20^, protein to mRNA ratios^20^ were derived from a study that investigated determinants of gene expression in several human tissues, including the heart; translation efficiency was computed using van Heesch’s RP^16^ and RNA-seq datasets; and finally, half-life values for mRNA^21^ and protein^22^ were obtained from studies with human cells (Fig 3). As controls, we used two other lists of genes derived from Eraslan’s study^20^: one containing the top 5% most abundant proteins in heart tissue and the other containing the 5% least expressed heart proteins^20^. As expected, the genes coding for the top 5% most abundant proteins had higher values for all analyzed parameters, compared to the overall to the genome (Fig 3A-F, compare blue bars with black bars, see also Table S2). Accordingly, the least expressed gene list had lower expression indicators, except for the protein half-life that was no different from the control group (Fig 3E). The high CV genes followed a particular profile, characterized by lower mRNA abundance, shorter half-lives (panel 3B, 3F), and higher TE (panel 3D). Combined, these features suggest that genes with high CV may be optimized for transient, yet fast and efficient expression.

**Figure 3.**
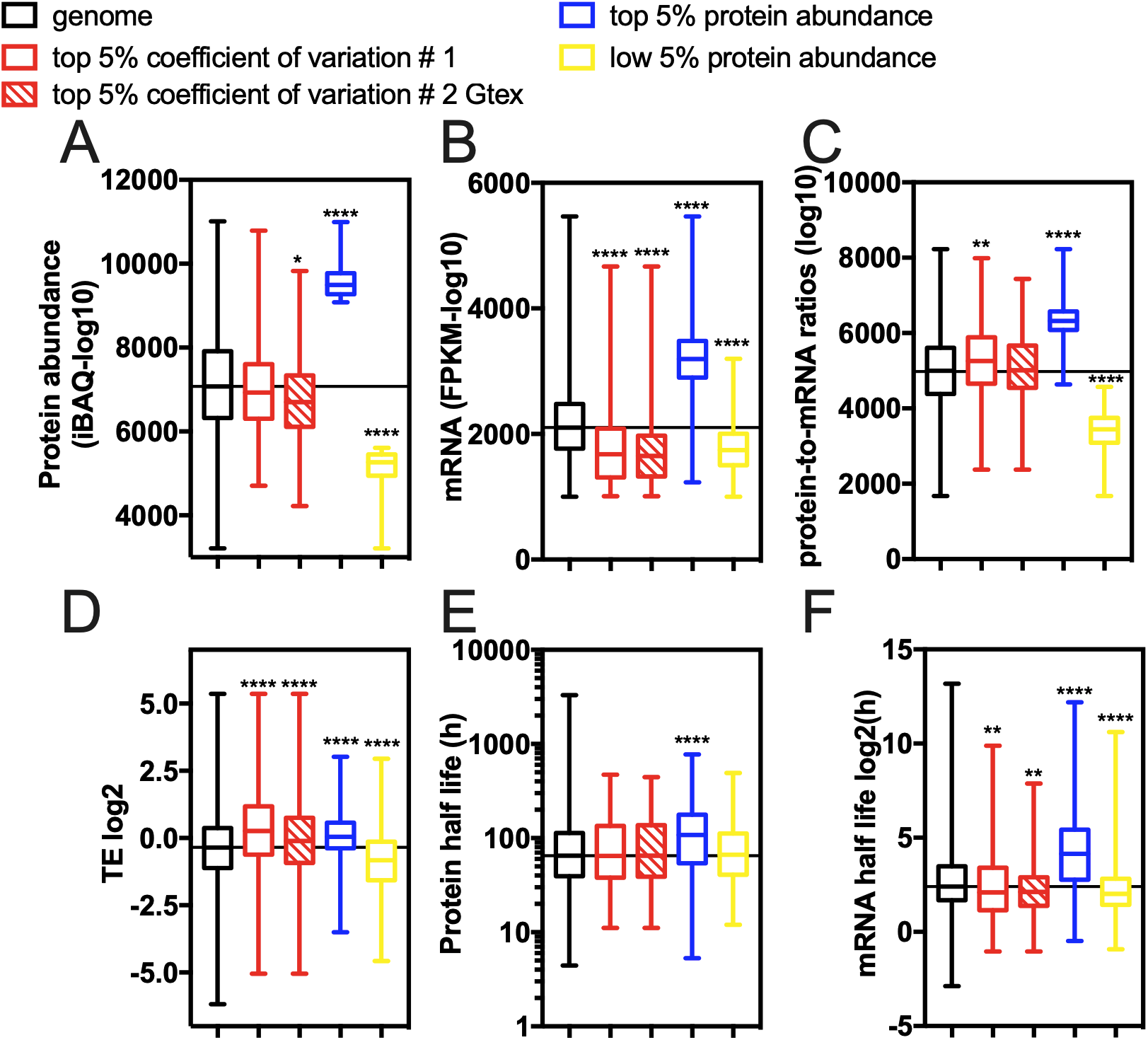
Indicators of expression for the genes with a high coefficient of variation (CV) in healthy human heart left ventricle tissue. (A) protein abundance^20^, (B) mRNA abundance^20^, (C) protein-to-mRNA ratios^20^, (D) translation efficiency (TE)^16^, (E) protein half-lives^22^, and (F) mRNA half-lives^21^ for genes with a high coefficient of variation of the genome. As a positive control, we used the top 5% most abundant proteins of cardiac proteome^20^. ANOVA Kruskal-Wallis test *0.011, **0.0054, and ****<0.0001.

The gene ontology analyzes (GO) of genes with high CV showed several pathways were over-represented, including immune response and metal homeostasis (Fig S2 and Table S3). Next, to identify co-regulation patterns within the high CV list #1, we used the mRNA-seq gene expression data for these 565 genes to build a correlation matrix (Fig 4A, 565 genes X 565 genes= 319,225 correlations). We defined eleven clusters. The gene interactions from each cluster were analyzed using the STRING database, which compiles protein-protein interactions (physical and functional)^23^. We found that 4 clusters were enriched in specific pathways Figure 4 B-E (see also Table S4). Genes related to the renal system process regulation were upregulated in individuals A, C, F, I, and L (Fig 4B). The other three main clusters were enriched in genes related to the immune system (Fig 4 D-F). Finally, we analyzed whether the genes with high CV in the cardiac transcriptome were previously associated with the disease. To address this point, we used DisGeNET, a platform that integrates disease-associated genes and variants from multiple sources^24^. The DisGeNET gives three kinds of scores: i) the “disease score” reflects the disease genes associations that are reported by several databases^24^; ii) the “disease specificity index” means that the gene or gene variant is disease-specific; and, iii) the “disease pleiotropy index” calculates the score for the disease-promiscuous genes or variants. The high CV genes had the same overall distribution of disease score values as the entire genome (Fig 5A, black bars compared with red bars). However, the high CV list had lower disease specificity index scores and higher scores for disease pleiotropy (Fig 5C). That is, genes with high CV among healthy individuals tend to be nonspecifically associated with the disease. Therefore, asymptomatic conditions or infections may account for much of the variation observed between individuals.

**Figure 4.**
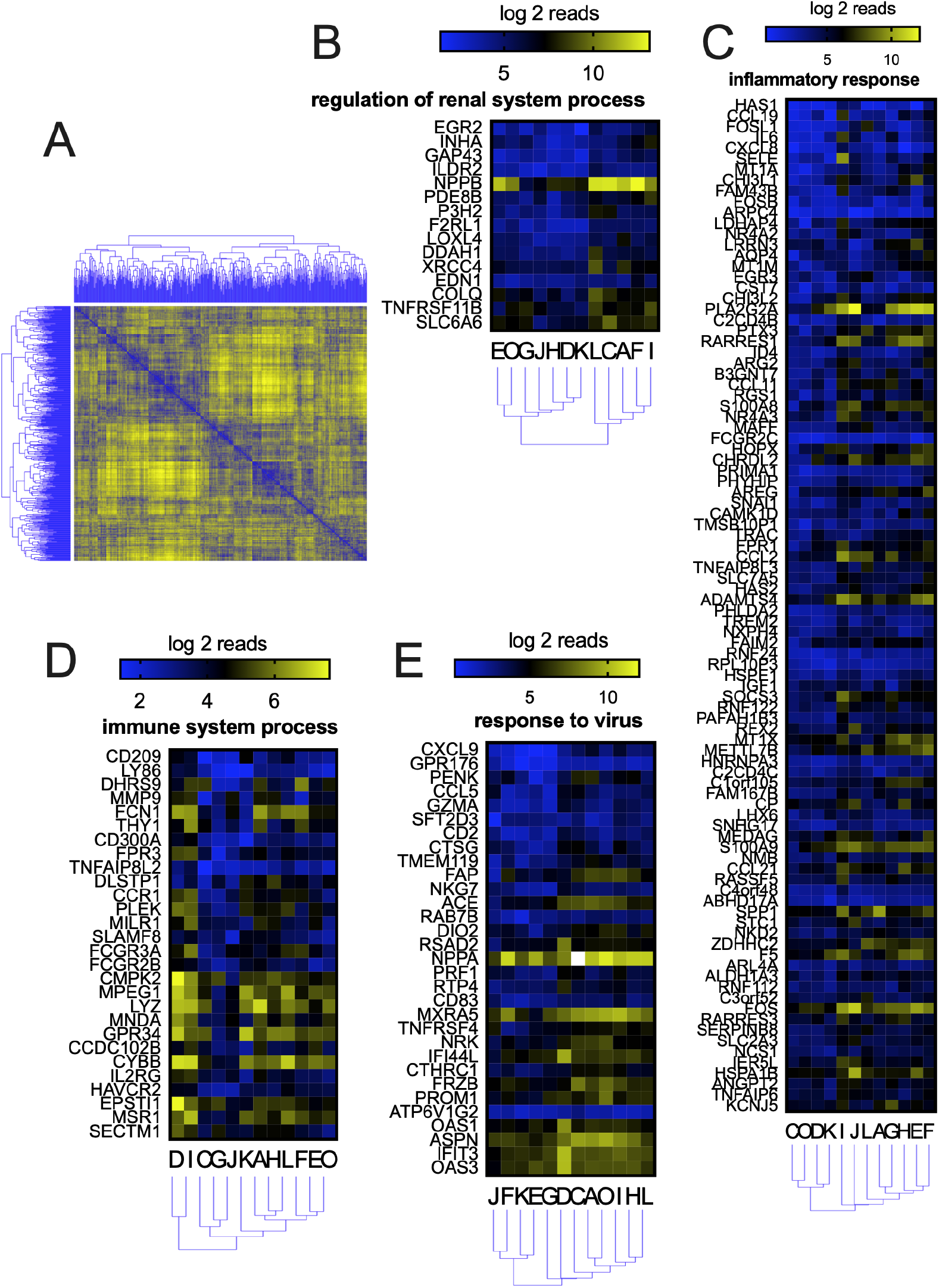
Co-regulation analyzes of genes with a high coefficient of variation (CV). (A) The distance matrix was created from the gene level (565 genes) clustering-based on co-regulation among twelve individuals. Eleven main clusters were found. Each cluster gene list was analyzed by STRING to found predicted protein-protein interactions (Table S3). Four main pathways were found, namely, regulation of renal system process (B), inflammatory response (C), immune system process (D), and response to the virus (E).

**Figure 5.**
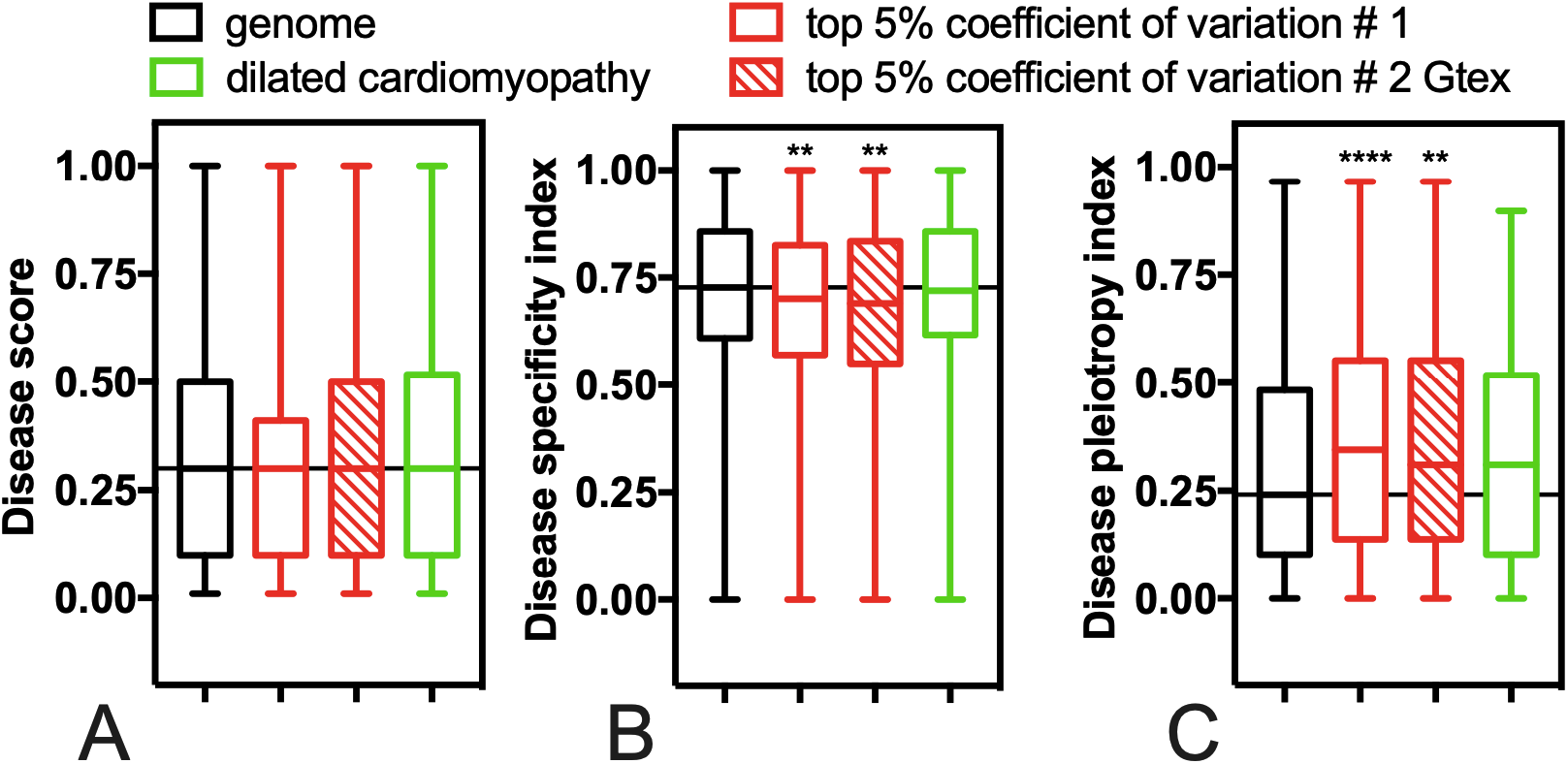
Genes with a high coefficient of variation are associated with pleiotropic diseases. The disease scores (A), disease specificity indexes (B), and disease pleiotropy indexes (C) were calculated for each gene using the DisGeNET database^24^. Boxplots compare the different disease indexes for genes with a high coefficient of variation (top 5%) with the genome. As a control, we used a list of genes identified as altered in dilated cardiomyopathy^16^. We expected to find differences among the dilated cardiomyopathy-related genes compared to the genome, but it was not the case. ANOVA Kruskal-Wallis test **0.006 and ****<0.0001.

We conclude that genes with a high coefficient of variation in cardiac transcriptome possess features associated with a high translational activity, such as high protein to RNA ratios and high translation efficiency (Fig 3C and 3D). A similar feature was observed with stress-related genes in yeast^25^. Since the expression of genes with high CV seems to respond to external stimulus (Fig 4), the translational features might be essential to a fast upregulation of adaptive pathways. Moreover, their short mRNA half-lives suggest transient regulation (Fig 3F). Interestingly, the genes with a high coefficient of variation were associated with several diseases (Fig 5C). This data can partially explain the differential susceptibility to diseases among human individuals, as previously described^26,27^.

## MATERIALS AND METHODS

| Data | Source |
| --- | --- |
| mRNA abundance (human heart) | van Heesch *et al.*, 2019 |
| mRNA abundance (human heart) | gtexportal.org/home/ |
| Ribosome profiling (human heart) | van Heesch *et al.*, 2019 |
| Dilated cardiomyopathy (DCM) genes | van Heesch *et al.*, 2019 |
| Translation efficiency (human heart) | van Heesch *et al.*, 2019 |
| Protein abundance (human heart) | Eraslan *et al.*, 2019 |
| mRNA abundance (human heart) | Eraslan *et al.*, 2019 |
| Protein per RNA amount (human heart) | Eraslan *et al.*, 2019 |
| Protein half-life (human hepatocyte) | Mathieson *et al.*, 2018 |
| mRNA half-life (human embryonic kidney 293 cells) | Narula *et al.*, 2019 |
| Disease score | Piñero *et al.*, 2020 |
| Disease specificity index | Piñero *et al.*, 2020 |
| Disease pleiotropy index | Piñero *et al.*, 2020 |

### Data sources

Coding sequences and annotation of *Homo sapiens* were obtained from the Ensembl Genome Browser (http://ensemblgenomes.org/).

### Statistical analyses, correlations, and raw data

The raw data used to create all Figures and the statistical analyses are presented in the Supplementary Tables (1-4). The statistical used for Figures 3 and 5 was the ANOVA Kruskal-Wallis test. All statistical analyses were performed with GraphPad Prism 7 software.

### GTEx data

To analyze the expression data from the GTEx database^3,4^ (https://gtexportal.org/home/) for the left ventricle, we wrote two algorithms in Python to get Left Ventricle experiments. The first uses the annotation file downloaded from the GTEx portal in .txt format and creates an output file containing only the annotations that contained the key term “left ventricle.” The second algorithm uses the output from GetTranscripts.py, and any RNA-seq data downloaded from the GTEx portal and writes as output only the data from the experiments listed in the GetTranscript.py output. The algorithms were deposited on Github repository https://github.com/RodolfoCarneiro/left_ventricle.

### STRING analyses

The STRING database (http://string-db.org/)^23^ was used to provide a critical assessment and integration of protein-protein interactions form the clusters of Figure 4.

### Clustering Analysis

All clustering analyses were performed by the Euclidean distance using Orange 3 software^28^.

### Translation efficiency calculation

Translation efficiency for each gene was calculated by dividing the RNA-seq reads by ribosome profiling reads. Next, the average translation efficiency for each gene from the 12 individuals was calculated (see Table S2).

### Calculation of coefficient of variation

The expression data were log-transformed (base = 2), and means, standard deviations, and gene to gene coefficients of variation were calculated. Genes with low expression were considered rare, i.e., those with an expression equal to or less than 1% of the genes in 75% (9 out of 12) of the samples. Similarly, those genes with high expression, i.e., those with an expression equal to or greater than 97.5% of the genes in 75% (9 out of 12) of the samples, were considered constitutive. After this, tests of equal coefficients of variation between each gene and the coefficient of variation of the mean expression of the genes selected as constitutive were calculated^29,30^. Genes with a higher coefficient of variation than the constituents were considered, those whose adjusted p-value by the Bonferroni method was less than 0.05 (alpha = 0.1) and whose coefficient of variation was higher than the coefficient of variation of the mean expression of constituent genes.

### Disease score database

The DisGeNET (http://disgenet.org/) was used as a platform to score the disease-associated genes and variants from multiple sources^24^.

## Supporting information

Support Information

## ACKNOWLEDGEMENT

We thank Rodrigo Requião and Carlos G. Schrago for helpful discussions. We thank Leonardo Maciel for critically reading the manuscript. We also thank Jared Nedzel from Broad Institute for help with the Gtex analyzes.

## CONFLICT OF INTEREST

The authors declare that they have no conflicts of interest with the contents of this article.

**Supplemental data for this article can be accessed on the publisher’s website.**

## REFERENCES

1. Hulse AM, Cai JJ. Genetic Variants Contribute to Gene Expression Variability in Humans. Genetics 2013; 193:95–108.

2. Frazer KA, Ballinger DG, Cox DR, Hinds DA, Stuve LL, Gibbs RA, Belmont JW, Boudreau A, Hardenbol P, Leal SM, et al. A second generation human haplotype map of over 3.1 million SNPs. Nature 2007; 449:851–61.

3. Aguet F, Brown AA, Castel SE, Davis JR, He Y, Jo B, Mohammadi P, Park Y, Parsana P, Segrè AV, et al. Genetic effects on gene expression across human tissues. Nature 2017; 550:204–13.

4. Consortium TGte. The GTEx Consortium atlas of genetic regulatory effects across human tissues. Science 2020; 369:1318–30.

5. Storey JD, Madeoy J, Strout JL, Wurfel M, Ronald J, Akey JM. Gene-Expression Variation Within and Among Human Populations. Am J Hum Genetics 2007; 80:502–9.

6. Heinig M, Adriaens ME, Schafer S, Deutekom HWM van, Lodder EM, Ware JS, Schneider V, Felkin LE, Creemers EE, Meder B, et al. Natural genetic variation of the cardiac transcriptome in nondiseased donors and patients with dilated cardiomyopathy. Genome Biol 2017; 18:170.

7. Heinig M. Using Gene Expression to Annotate Cardiovascular GWAS Loci. Frontiers Cardiovasc Medicine 2018; 5:59.

8. Stranger BE, Nica AC, Forrest MS, Dimas A, Bird CP, Beazley C, Ingle CE, Dunning M, Flicek P, Koller D, et al. Population genomics of human gene expression. Nat Genet 2007; 39:1217–24.

9. Stranger BE, Forrest MS, Dunning M, Ingle CE, Beazley C, Thorne N, Redon R, Bird CP, Grassi A de, Lee C, et al. Relative Impact of Nucleotide and Copy Number Variation on Gene Expression Phenotypes. Science 2007; 315:848–53.

10. Spielman RS, Bastone LA, Burdick JT, Morley M, Ewens WJ, Cheung VG. Common genetic variants account for differences in gene expression among ethnic groups. Nat Genet 2007; 39:226–31.

11. Crawford DL, Oleksiak MF. The biological importance of measuring individual variation. J Exp Biol 2007; 210:1613–21.

12. Whitehead A, Crawford DL. Variation in tissue-specific gene expression among natural populations. Genome Biol 2005; 6:R13.

13. Oleksiak MF, Roach JL, Crawford DL. Natural variation in cardiac metabolism and gene expression in Fundulus heteroclitus. Nat Genet 2005; 37:67–72.

14. Ingolia NT, Brar GA, Rouskin S, McGeachy AM, Weissman JS. The ribosome profiling strategy for monitoring translation in vivo by deep sequencing of ribosome-protected mRNA fragments. Nat Protoc 2012; 7:1534–50.

15. Cenik C, Cenik ES, Byeon GW, Grubert F, Candille SI, Spacek D, Alsallakh B, Tilgner H, Araya CL, Tang H, et al. Integrative analysis of RNA, translation, and protein levels reveals distinct regulatory variation across humans. Genome Res 2015; 25:1610–21.

16. Heesch S van, Witte F, Schneider-Lunitz V, Schulz JF, Adami E, Faber AB, Kirchner M, Maatz H, Blachut S, Sandmann C-L, et al. The Translational Landscape of the Human Heart. Cell 2019; 178:242–260.e29.

17. Ingolia NT, Ghaemmaghami S, Newman JRS, Weissman JS. Genome-Wide Analysis in Vivo of Translation with Nucleotide Resolution Using Ribosome Profiling. Science 2009; 324:218–23.

18. Weinberg DE, Shah P, Eichhorn SW, Hussmann JA, Plotkin JB, Bartel DP. Improved Ribosome-Footprint and mRNA Measurements Provide Insights into Dynamics and Regulation of Yeast Translation. Cell Reports 2016; 14:1787–99.

19. Fang H, Huang Y-F, Radhakrishnan A, Siepel A, Lyon GJ, Schatz MC. Scikit-ribo Enables Accurate Estimation and Robust Modeling of Translation Dynamics at Codon Resolution. Cell Syst 2018; 6:180–191.e4.

20. Eraslan B, Wang D, Gusic M, Prokisch H, Hallström BM, Uhlén M, Asplund A, Pontén F, Wieland T, Hopf T, et al. Quantification and discovery of sequence determinants of protein-per-mRNA amount in 29 human tissues. Mol Syst Biol 2019; 15:e8513.

21. Narula A, Ellis J, Taliaferro JM, Rissland OS. Coding regions affect mRNA stability in human cells. Rna 2019; 25:1751–64.

22. Mathieson T, Franken H, Kosinski J, Kurzawa N, Zinn N, Sweetman G, Poeckel D, Ratnu VS, Schramm M, Becher I, et al. Systematic analysis of protein turnover in primary cells. Nat Commun 2018; 9:689.

23. Szklarczyk D, Gable AL, Lyon D, Junge A, Wyder S, Huerta-Cepas J, Simonovic M, Doncheva NT, Morris JH, Bork P, et al. STRING v11: protein–protein association networks with increased coverage, supporting functional discovery in genome-wide experimental datasets. Nucleic Acids Res 2018; 47:gky1131.

24. Piñero J, Ramírez-Anguita JM, Saüch-Pitarch J, Ronzano F, Centeno E, Sanz F, Furlong LI. The DisGeNET knowledge platform for disease genomics: 2019 update. Nucleic Acids Res 2019; 48:D845–55.

25. Carneiro RL, Requião RD, Rossetto S, Domitrovic T, Palhano FL. Codon stabilization coefficient as a metric to gain insights into mRNA stability and codon bias and their relationships with translation. Nucleic Acids Res 2019; 47:2216–28.

26. Li J, Liu Y, Kim T, Min R, Zhang Z. Gene Expression Variability within and between Human Populations and Implications toward Disease Susceptibility. Plos Comput Biol 2010; 6:e1000910.

27. Wegler C, Ölander M, Wiśniewski JR, Lundquist P, Zettl K, Åsberg A, Hjelmesæth J, Andersson TB, Artursson P. Global variability analysis of mRNA and protein concentrations across and within human tissues. Nar Genom Bioinform 2019; 2.

28. Demšar J, Curk T, Erjavec A, Machine ČGTJO. Orange: Data Mining Toolbox in Python. J Mach Learn Res 2013; 14:2349–53.

29. Feltz CJ, Miller GE. An Asymptotic test for the equality of coefficients of variation from k populations. Stat Med 1996; 15:647–58.

30. Krishnamoorthy K, Lee M. Improved tests for the equality of normal coefficients of variation. Computation Stat 2014; 29:215–32.

