## Supplementary material for "Natural variation of the cardiac transcriptome in humans": Support Information

### Support information for:

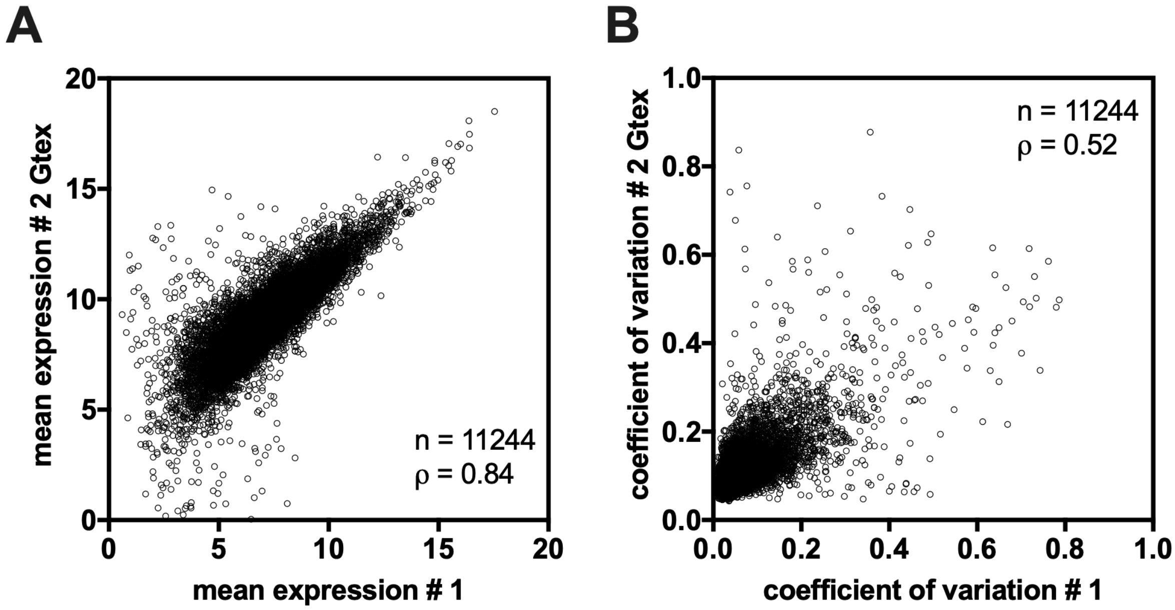


**Figure S1. Correlation of mean of expression of mRNA (A) and coefficient variation (B) for each gene among the cohort from van Heesch's study (#1) and Gtex (#2 Gtex).** The number of genes (n) was 11224 for both analyzes while the Spearman correlation coefficient ($\rho$) were 0.84 and 0.52 for A and B respectively.

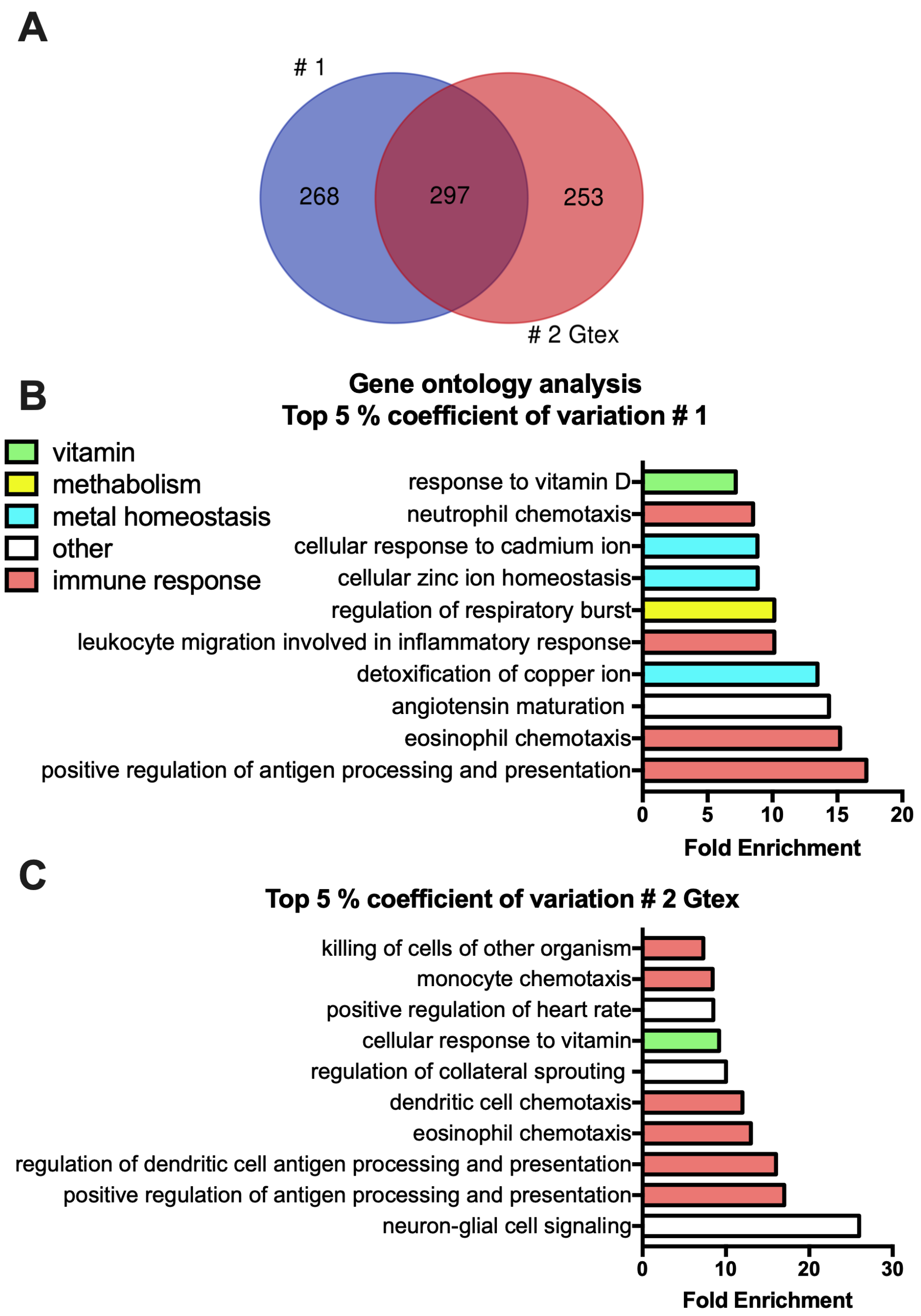


**Figure S2. Gene ontology of the top 5% genes with higher coefficient of variation from van Heesch's study (#1) and Gtex (#2 Gtex).** (A) Venn diagram showing that more than 50% of the top 5% genes with higher coefficient of variation from van Heesch's study (#1) and Gtex (#2 Gtex) are the same. Gene ontology of top 5% genes with higher coefficient of variation from van Heesch's study (B) and Gtex (C). All analyzes were derived of mRNA-seq data from human heart left ventricle using 12 (#1) or 432 (#2 Gtex) healthy donors.
